# On the role of photoreceptor identity in controlling accurate wiring of the *Drosophila* visual circuit

**DOI:** 10.1101/2020.10.13.337865

**Authors:** Weiyue Ji, Lani F. Wu, Steven J. Altschuler

## Abstract

During development, neurons extend in search of synaptic partners. Precise control of axon extension velocity can therefore be crucial to ensuring proper circuit formation. How velocity is regulated – particularly by the extending axons themselves – remains poorly understood. Here, we investigate this question in the *Drosophila* visual system, where photoreceptors make precise connections with a specific set of synaptic partners that together create a circuit underpinning neural superposition (NSP). We used a combination of genetic perturbations and quantitative image analysis to investigate the influence of cell identity on growth cone velocity and subsequent spatial-temporal coincidence of presynaptic and postsynaptic neurons. Our study provides a case study of how cell autonomous properties of presynaptic axons play a pivotal role in controlling the dynamics of growing axons and determining the formation of a precise neuronal circuit.

## Introduction

One of the most fascinating questions in neuroscience is how complex neural circuits form. During development, neurons choose synaptic partners within brief time intervals. How do they arrive at the right place and at the right time? One way to ensure proper circuit formation is through precise control of extending growth cone velocities. Previous studies have shown that regulation of velocity can be achieved through external signals, such as neurotransmitters and morphogens (Castellani, 2013; Petros et al., 2008). However, the degree to which neurons autonomously regulate their velocity is less well understood.

The *Drosophila* visual system offers a remarkable opportunity to investigate cell-autonomous control of growth cone extension. Each eye comprises 800 unit eyes, or ommatidia, and each ommatidium contains 8 photoreceptors (“R cells”, labeled R1 through R8). Axons of the photoreceptors extend from the eye to the optic lobe of *Drosophila* in groups of 800 “bundles” with canonical intra-bundle orientations. Six of the eight R cells (R1-R6) stop extending at the lamina plexus and wire to their postsynaptic targets, the lamina monopolar cells (“L cells”). At each bundle, the R1-R6 cells diverge towards six different L-cell targets; and, at each L-cell target, R1-R6 cells converge, each originating from a different bundle (Figure S1A). This final complex, yet highly stereotyped neuronal circuit topology is referred to as neural superposition (NSP) (Agi et al., 2014; Ting & Lee, 2007).

An intriguing discovery is that R1-R6 each exhibit relatively consistent and cell subtype-specific velocity (i.e. direction and speed) during growth cone extension (Langen et al., 2015). This suggests that the velocity of growth cones plays a pivotal role in their synaptic partner selection. Previous studies suggested that both ommatidia orientation of the originating bundle (Clandinin & Zipursky, 2000) and interactions among growth cones within the same bundle (Chen & Clandinin, 2008; Schwabe et al., 2013) contribute to the direction of R cell projections. However, no studies have investigated the role of a photoreceptor’s identity in controlling its own growth cone extension.

In this study, we used a combination of genetic perturbations and quantitative image analysis to study the role of cell identity in controlling accurate wiring of the *Drosophila* visual circuit. First, we utilized the local regularity of the circuit to develop a data-driven standardized coordinate system around each bundle, which facilitated comparisons across samples. Then, we quantified extension dynamics of wild-type photoreceptor growth cones across different developmental stages. Next, we investigated how these dynamics change when cell fates of growth cones are altered under genetic perturbations. Finally, we estimated the impact of nearest neighbor interactions in determining extension velocity. Together, our quantitative study revealed that photoreceptor identities determine their speed but not angle of extension, and that differences in extension speed ultimately contribute to target specificity. Our study provides a case study of cell autonomous mechanisms in neuronal circuit formation and highlights the importance of disentangling the regulation of speed and direction – the two components of velocity – in developmental processes.

## Results

### Changes in R3 or R4 cell identity lead to changes in final targeting

The role of photoreceptor identity during development has been extensively studied. Photoreceptors develop in three sequential pairs during eye development: R2/R5, then R3/R4 and last R1/R6 (Nériec & Desplan, 2016; Treisman, 2013). The R3/R4 pair is particularly important, as the asymmetry between R3 and R4 determines the 90-degree rotation of R-cell clusters in the developing eye disc and later, in the asymmetric trapezoidal arrangement of the adult ommatidia (Treisman, 2013). Interestingly, the R3/R4 pair also stands out in the wiring diagram of NSP. The target positions of R3 and R4 (“T3” and “T4”, respectively) are asymmetrical, while the target positions of the other two pairs (R1/R6, R2/R5) are symmetrical (Figure 1 – figure supplement 1, A). Therefore, we focused our effort on understanding the role of R3/R4 identities—and their contribution to asymmetric targeting—during the NSP wiring process in the lamina.

To alter R3 and R4 cell identities, we used genetic perturbations in the planar-cell-polarity pathway. Specifically, over-expression of Frizzled (*Fz*) with *sevenless* (*sev*) enhancer (*sev>Fz*) generates ommatidia with two R3s (Strutt et al., 1997; Zheng et al., 1995), while over-expression of the intracellular domain of Notch (*N*^*ic*^) under the same enhancer (*sev>N*^*ic*^) generates ommatidia with two R4s (Cooper & Bray, 1999; Fanto & Mlodzik, 1999). To visualize the perturbed bundles, we used a membrane-bound red fluorescent protein (RFP) under the same enhancer. Since the lamina plexus is densely packed with photoreceptors, it is challenging to disambiguate individual photoreceptor growth cones. Thus, we induced our perturbation sparsely. We utilized a FRT-dependent GAL80 “flip-in” construct together with a heat-shock activated flippase to generate sparse clones of perturbed bundles (Bohm et al., 2010; Chou & Perrimon, 1996). Further, to differentiate between R3s and R4s cell types at an early stage of development in both the ommatidia and lamina, we utilized the R4-specific enhancer *mδ05* fused with a membrane GFP protein (*mδ-GFP*) (Cooper & Bray, 1999).

We visually inspected specimens of sparsely perturbed *sev>Fz* and *sev>N*^*ic*^ flies after the completion of NSP wiring (> 36 hours after puparium formation, or hrs APF; see Figure 1 – figure supplement 1, B for timeline (Agi et al., 2014)). We found that changes in cell identity resulted in changes in final targeting (Figures 1, Figure 1 – figure supplement 2). For bundles with two R4 photoreceptors, both the normal and fate-transformed R4s target the canonical R4 target, T4. For bundles with two R3 photoreceptors, the normal R3 targets the canonical R3 target, T3, while the fate-transformed R3 targets T3’, a new target position that is mirror-symmetric to T3, instead of the original T3. (See Supplementary file 1A for phenotype penetrance.) This result raised the hypothesis that cell identity interacts with bundle position to shape the targeting of R3 and R4 photoreceptors.

**Figure 1.**
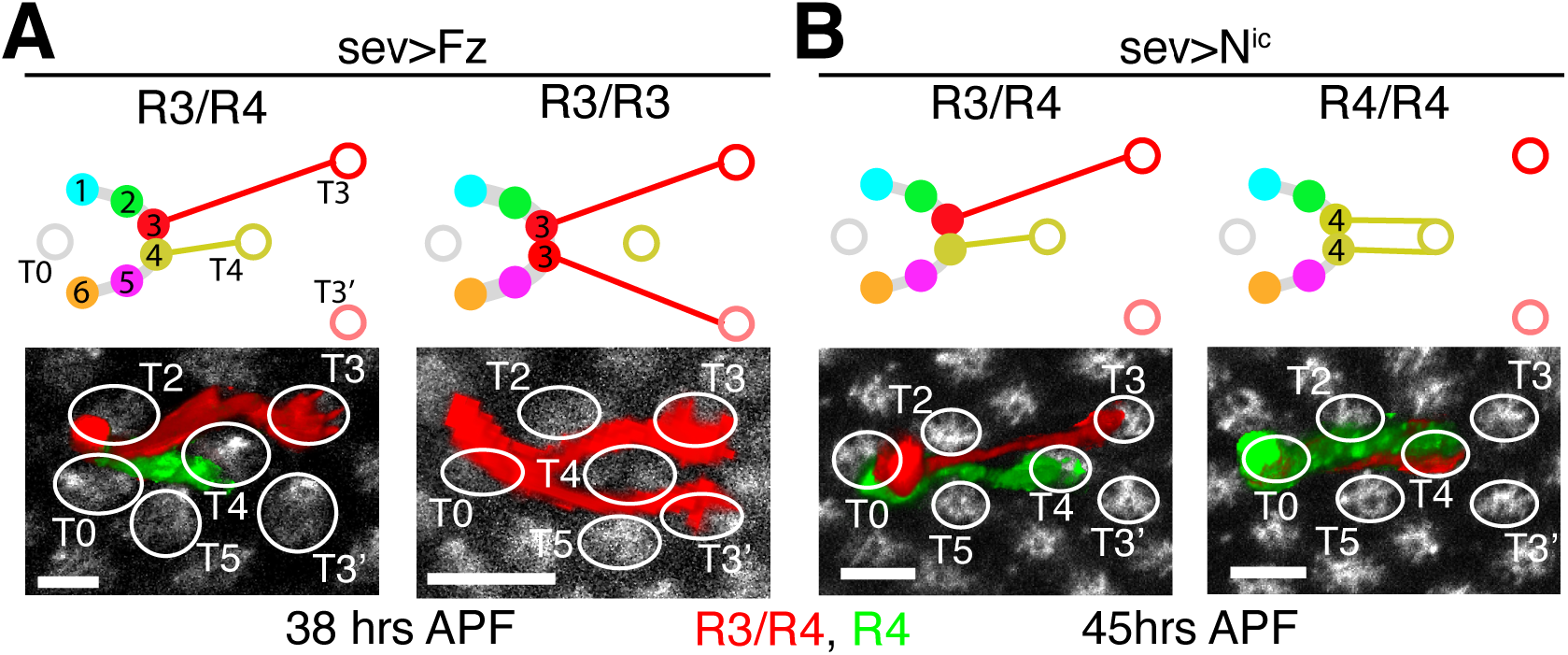
Changes in R3 or R4 cell identity lead to changes in final targeting. **(A-B)** Representative schematics and images of bundles from (A) *sev>Fz* (38 hrs APF) and (B) *sev>N*^*ic*^ (45 hrs APF) flies (see Figure 1 – figure supplement 2 for more examples). Top panels: schematic of wild-type or altered wiring topology. Solid or open circles: starting points (‘heels”) or targets (respectively); colors coordinated between R cells and targets. T3’: target of fate-altered R3s; T0: target located within the bundle of interest (though targeted by R cells from other bundles in NSP wiring). Bottom panels: confocal images of representative bundles. Photoreceptor growth cones were segmented, pseudo-colored, and intensity scaled for visualization (Methods). Red: *sev>RFP* expression; green: *mδ-GFP* expression; white: Fasciclin 2 (FasII) antibody staining. White ellipses: targets. Scale bar: 5 μm.

### Wild-type R3s and R4s exhibit asymmetric speeds but symmetric directions of extension

Perturbing cell identity changed the final target choice. But, did it also change the behavior of early extension? We next focused on quantitative characterization of growth cone extension during NSP.

Characterizing patterns in growth cone extension requires comparing growth cone morphology across multiple bundles and between animals. The wiring of the *Drosophila* lamina exhibits a crystal-like two-dimensional pattern for the 800 bundles across the lamina plexus. However, variations in image orientation and irregular local warping of the lamina plexus confounds bundle-to-bundle and sample-to-sample comparison. We therefore developed an approach that utilizes the local regularity of a growth cone’s environment and quantifies its behavior according to a standardized polar coordinate system (Figure 2; Methods). For each bundle, we identified the starting positions of all R-cells (Heel grid, Figure 2A) and their putative targets (Target grid, Figure 2B). We also identified a center point “C”, which lies at the intersection of the line connecting R3 and T3 and the line connecting R4 and T3’. We then extrapolated polar coordinates by normalizing (Figure 2C): 1) length, so that |C-T4| = 1 (A.U.); and 2) angle, so that ∡(T3,C,T4) = ∡(T4,C,T3’) = 1 (A.U.) and that T3 and T3’ were placed at angles +1 and −1, respectively. (We note that in our samples, |C-T4| is 7.3 ± 1.5 *μ*m and ∡(T3,C,T3’) is 27.8 ± 6.8°.)

**Figure 2.**
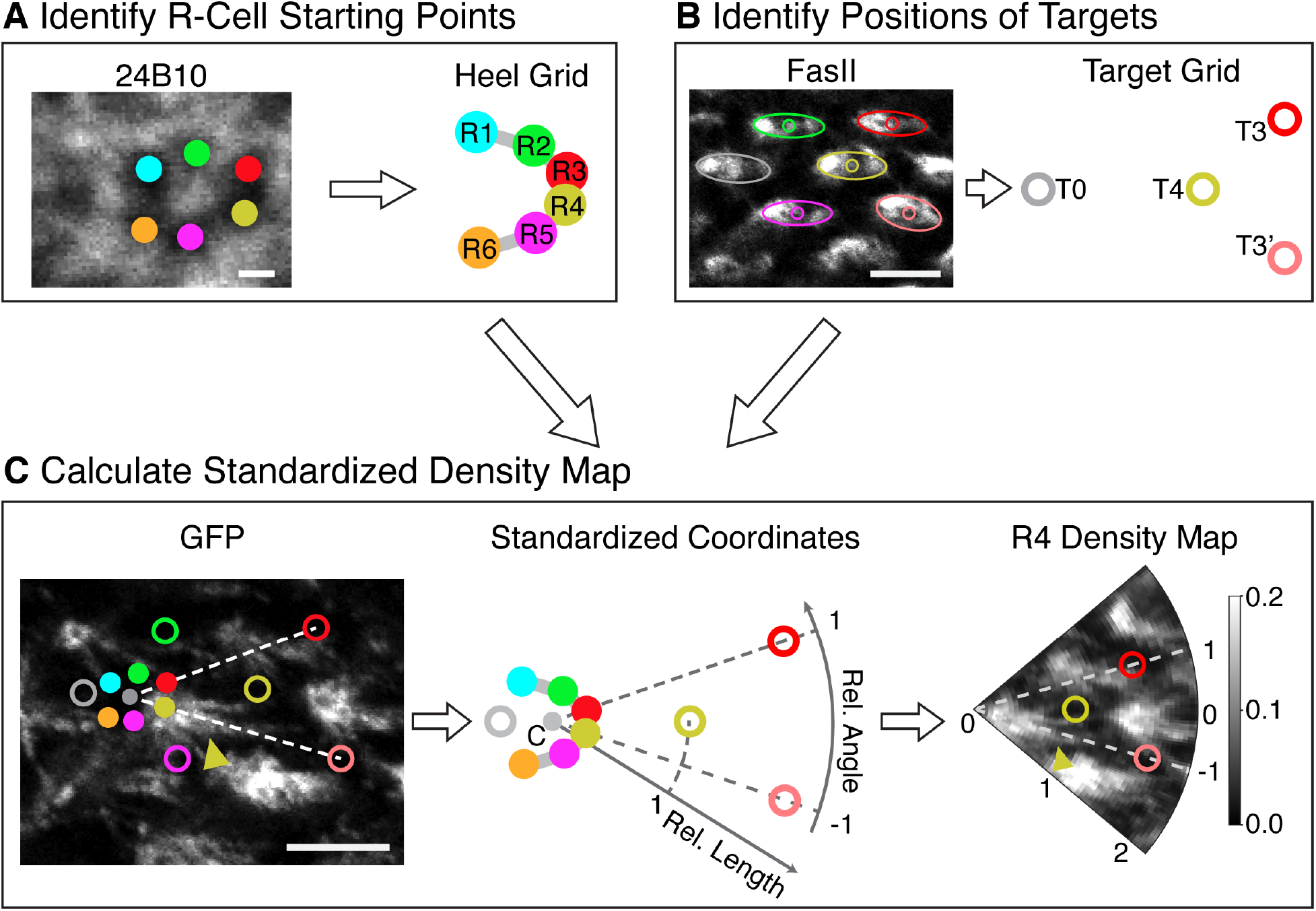
Establishment of standardized coordinates for comparison of growth cone extension. Overview of the image quantification process. **(A)** The heel grids (i.e. R-cell starting points) are identified (via 24B10 antibody labeling R-cell membranes; Methods). **(B)** The target grid is identified (via FasII antibody labeling L-cells membranes; Methods). **(C)** In each region, (left) the annotation (from A-B) is used (middle) to produce standardized coordinates that are used (right) to transform confocal images into standardized density maps. This transformation allows relative lengths (radial coordinate) and angles (angular coordinate) of extending growth cones to be compared across regions. Images: confocal images of a bundle region were sampled from one wild-type fly at 26 hrs APF. Images were cropped, re-oriented and intensity scaled for visualization (Methods). Scale bars: 5 μm for FasII and GFP images; 1 μm for 24B10 image.

Using these standardized coordinates, we examined behavior of wild-type flies taken from early to late stage of neural superposition (22 hrs to 36 hrs) to characterize the dynamics of R3 and R4 growth cones during extension. First, we examined changes in the morphologies of R3 and R4 growth cones over time. By 22 hrs APF, growth cones of both R3s and R4s have already polarized (Figures 3A and Figure 3 – figure supplement 1), with numerous filopodia at the leading edge. This morphology is maintained until ~32 hrs APF, after which the number of filopodia decreases and the growth cones take on narrower, more linear morphologies (Figure 3A).

**Figure 3.**
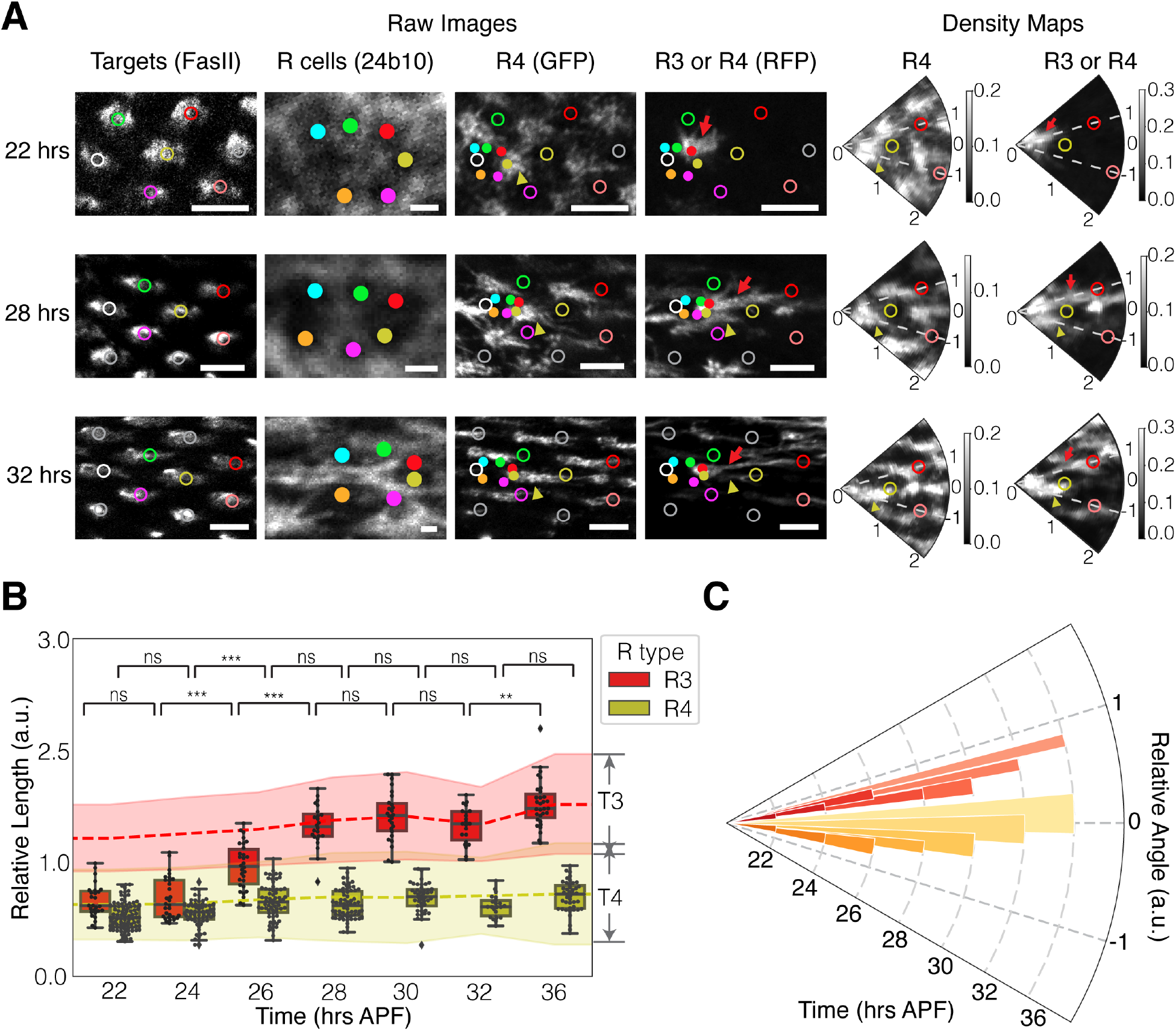
Wild-type R3s and R4s exhibit asymmetric speeds but symmetric directions of extension. **(A)** Representative images of bundles from wild-type flies at 22, 28, and 32 hrs APF (See Figure 3 - figure supplement 1 for examples at different timepoints). Left: Confocal images of example bundles. From left to right: FasII channel labeling the target cells; 24B10 channel labeling membrane of all R cells; GFP channel labeling membrane of R4 cells; RFP channel labeling membrane of R3 or R4 cells. Right: Density maps of GFP (R4 cells) and RFP channel (R3 or R4 cells) after coordinate transformation. Images were cropped, re-oriented and intensity scaled individually for visualization (Methods). R-cells and targets indicated and colored as in Figure 1; white circles: T0; gray circles: other targets. Yellow arrowheads: R4 growth cones; red arrows: R3 growth cones. Scale bars: 5 μm for FasII, GFP and RFP images; 1 μm for 24B10 images. **(B)** Change in relative lengths of wild-type R3 and R4 growth cones over time. Red and yellow dashed lines: mean of the center of target positions for T3 and T4, respectively. Error bars of the dashed lines: mean of the upper and lower boundaries of targets. Significance: calculated using a two-sided Mann-Whitney test with p values adjusted by Bonferroni method. ns: p > 0.5, *: 0.01 < p < 0.05, **: 0.001 < p < 0.01, ***: p < 0.001. Sample sizes (number of bundles) of each time point: R3 growth cones (n = 27, 30, 30, 24, 26, 22, 31); R4 growth cones (n = 96, 67, 67, 67, 44, 22, 43). n ≥ 2 biological replicates for each time point. Supplementary file 1B for p-values. **(C)** Change in relative angles of wild-type R3 and R4 growth cones over time. Red and yellow bars represent R3s and R4s, respectively. Plotted bars: radial coordinate indicates time; angle coordinate indicates mean relative angle of R3 or R4 growth cones at the given time point; and width indicates standard deviation of angle values at the given time point. Sample sizes as in (B). See Supplementary file 1B for p-values.

Next, we examined the speed and angle of growth cones and their relationship with their targets. The growth cone of R4 is close to T4 as early as 22 hrs APF, with some of the filopodia at the leading edge of the growth cone already appearing to make contact, and fully reach T4 by 28 hrs APF. R3 growth cones are initially similar in length to R4 at 22 hrs APF, yet they continue to extend past T4 starting from 24 hrs APF and reach the more distant target T3, by 28 hrs APF (Figure 3B). Thus, R3s have a markedly higher speed of extension than R4s — a roughly 8-fold difference (0.16 a.u./hour vs. 0.02 a.u./hour). While the speed of R3 and R4 are asymmetric, their extension angles are, surprisingly, symmetrical at early stages of NSP wiring (Figure 3C). We observed dramatic changes in the extension angle of R4s at 32 hrs APF and R3s at 36 hrs APF, correlating with morphological changes that indicate the transition from axon pathfinding to synapse formation.

### Extension speed is responsible for asymmetrical targeting

Based on these observations in wild-type, we hypothesized that extension speed is responsible for the asymmetrical targeting of R3/R4 pairs. To test this hypothesis, we examined the behavior of sparsely perturbed mutant (*sev>Fz* and *sev>N*^*ic*^) flies that have both wild-type-like and mutant bundles at 24 or 28 hrs APF, corresponding to early or late stages of their extension (Figures 4 and Figure 4 – figure supplement 1).

**Figure 4.**
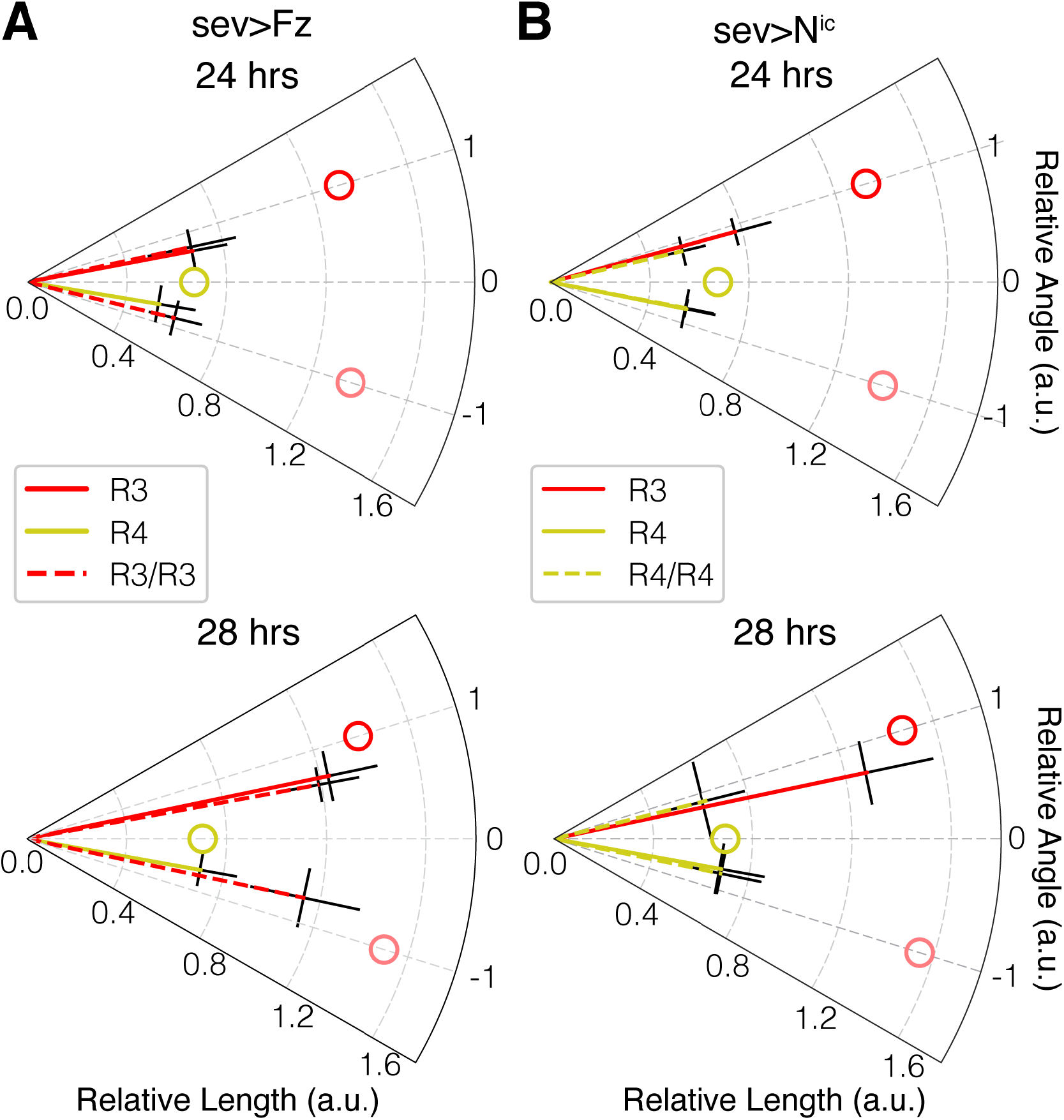
Extension speed is responsible for asymmetrical targeting. Polar plots of relative lengths and angles for wild-type-like (R3/R4) and perturbed (R3/R3 or R4/R4) bundles under **(A)** *sev>Fz* and **(B)** *sev>N*^*ic*^ conditions at 24 and 28 hrs APF. See Figure 4 – figure supplement 1 for representative images of each condition. Radial coordinate: relative length; angle coordinate: relative angle; error bars: standard deviations for length or angle. Solid lines: control bundles; dashed lines: perturbed bundles. Open circles: mean of the center of target positions. Sample numbers for growth cones at wild-type R3 position, wild-type R4 position, perturbed R3 position, and perturbed R4 position, respectively: *sev>Fz* at 24 hrs (n = 24, 38, 10, 10) or 28 hrs (n = 52, 151, 23, 23); *sev>N*^*ic*^ at 24 hrs (n = 23, 46, 10, 10) or 28 hrs (n = 39, 113, 29, 29). n ≥ 3 biological replicates for each genetics and time point. Supplementary file 1C for p-values.

We compared the behaviors of fate-transformed to wild-type bundles. The fate-transformed bundles contain either two R3s or two R4s (one of which is fate transformed and the other wild-type). We observed that the extension speed and angle of the wild-type cells in the fate-transformed bundles is similar to the R cells of the same identity in the wild-type-like bundles. More importantly, we discovered that within the fate-transformed bundles, the two R cells extend at similar speeds and symmetric angles (Figure 4). Thus, we conclude that cell identity is instrumental in determining extension speed of photoreceptor growth cones, which ultimately lead to the asymmetrical targeting of R3 and R4s.

Taken together, the initial directions of extension for R-cells at the R3 and R4 positions are mirror symmetric and independent of cell identity (i.e., whether wild-type or fate-transformed). What could determine the initial extension directions? Simulations based on the geometric configurations of R-cell bundles and their targeting positions (Langen et al., 2015) suggested the possibility of a simple repulsion rule. That is, equal repulsion from immediate neighbor(s) within the same bundle determines the extension direction of an R cell. While this appeared reasonable for most of the R cells, R3/4 have asymmetrical targets and it was unclear how this rule could apply to their initial extension directions. In light of the discovery that R3 and R4 initially extend symmetrically, we re-examined the possibility of a simple repulsion rule (Figure 4 – figure supplement 2, A). Analysis showed that this would lead to predicted extension angles that are ~30° further from the bundle midline than those that are experimentally measured (Figure 4 – figure supplement 2, B). Instead, regression analysis suggested that R2 or R5 contribute ~2x the repulsion of the other immediate neighbor R3 or R4 (e.g. for the case of R3, R2 has twice the repulsion of R4) (Figure 4 - figure supplement 2, B). Thus, our quantitative study of growth cone extension suggests the hypothesis that a weighted repulsion model among neighbors within a bundle determines extension directions during patterning of the NSP circuit.

## Discussion

The velocity, defined by both direction and speed of the growth cone, is an important property of extending neurons during circuit formation. How is velocity controlled during neuronal development, and what are the consequences of changes in velocity in vivo for circuit formation? Here, we investigate this question in the context of *Drosophila* NSP circuit, in which ~4800 neurons (= 800 bundles x 6 R-cell types) swap relative positions and identify their targets with astonishing accuracy (Horridge & Meinertzhagen, 1970). To compare the dynamics across populations of neurons, we developed a standardized coordinate system for describing growth cone position. This coordinate system was essential in overcoming bundle-to-bundle and fly-to-fly heterogeneity, and similar approaches could be adapted to quantify and compare the dynamics of neurons in other developmental systems with other wiring geometries. We found that cell identities for R3 and R4 neurons determine their speed but not direction of extension. As a consequence, while R3 and R4 start with symmetric extension directions, their differences in speed lead to extension length differences by the final time of wiring and subsequently to asymmetric target choices. These observations highlight a crucial role for cell-autonomous mechanisms in controlling the dynamics of neuronal extension and, ultimately, the spatial-temporal coincidence of presynaptic and postsynaptic neurons.

One simple self-organizing mechanism to determine extension direction of photoreceptor growth cones is through repulsion of neighboring growth cones within the same bundle. We discovered that a simple repulsion model—where each R cell contributes equally—is not sufficient to explain the measured extension direction; rather, a model is supported in which R2/5 contributes much more strongly than R3/4. Such differential repulsion could arise through different strengths of expressed repulsive cues among neighboring cells. Interestingly, previous studies did show that Flamingo, a seven-pass transmembrane cadherin capable of inducing repulsion in both axons and dendrites (Matsubara et al., 2011; Senti et al., 2003), is differentially expressed amongst the different R-cell types, with R2/R5 having ~3x the expression levels of R3/R4 (Schwabe et al., 2013). Our study provides a quantitative framework to investigate molecular mechanisms underlying this developmental decision process.

Our study also provided a functional readout and framework for future investigations into mechanisms of how cell identity controls growth cone dynamics. Regulation of cytoskeletal dynamics is a clear possibility. Studies in cultured mammalian neurons have identified key components of the cytoskeleton that regulate axon outgrowth and how these components can be regulated by signaling molecules (Dent et al., 2011; Geraldo & Gordon-Weeks, 2009). For example, the cell-surface receptor Notch has been shown to regulate the speed of neurite outgrowth (Berezovska et al., 1999; Franklin et al., 1999; Šestan et al., 1999) via controlling the stability of microtubules (G. Ferrari-Toninelli et al., 2008) and the expression levels of signaling proteins (Giulia Ferrari-Toninelli et al., 2009). In the *Drosophila* NSP circuit, there is evidence that cytoskeletal regulators, such as Genghis khan and Cdc42, plays a crucial role in precise targeting of photoreceptors (Gontang et al., 2011). Additionally, while we focused on extension velocity, our findings do not exclude other factors that may also contribute to growth cone wiring that are also changed when cell identity is transformed. Understanding how cell identities translate to differences in molecular profiles and finally to changes in the cytoskeletal networks that control growth cone dynamics, will provide valuable insight into how growth cone velocity can be controlled during neuronal circuit development *in vivo*.

Changes in neurite speed have been observed in multiple models of neuronal circuit formation. For example, axons that require midline crossing often exhibit large changes in growth cone kinetics adjacent to the midline. Such pausing behavior is observed in crossing of mice retinal axons (Godement et al., 1994; Mason & Wang, 1997), zebrafish retinal ganglion cells (Hutson & Chien, 2002), and zebrafish postoptic commissures (Bak & Fraser, 2003). Growth cones of Zebrafish Mauthner cells also pause briefly when encountering successive motor neurons while extending along the midline (Jontes et al., 2000). There are also examples of changes in neurite velocity that have functional consequences both *in vitro* and *in vivo*. In studies of cultured snail *Helisoma* neurons, it has been shown that changes in ganglion neuronal speed correlate with their ability to form electrical synapses among themselves (P. Haydon et al., 1984; P. G. Haydon et al., 1987). Furthermore, there is evidence in both vertebrate and invertebrate sensory circuits that direction and speed of axon-dendritic interactions can contribute to connection specificity (Balaskas et al., 2019; Kiral et al., 2020). Our study in the *Drosophila* neural superposition demonstrates how cell-autonomous control of velocity is instrumental in the precise wiring of a complex sensory circuit. Taken together, these studies highlight the importance of velocity and vector-based strategies in developmental neuroscience.

## Materials and methods

### Fly stocks and handling

Fly stocks were constructed and maintained at 25 °C using standard protocols.

The following fly lines were used in this study: tubP>stop>GAL80 (II), tubP>stop>GAL80 (III) and hs-FLP122 (X) (gifts from T. Clandinin, Stanford University), mδ05-CD4::GFP (III) (gift from P.R. Hiesinger, Free University Berlin), Uas-N^ic^ (II) (gift from B.A. Hassan, ICM Institute for Brain and Spinal Cord); UAS-CD4::tdTomato (II) (gift from L.Y. Jan and Y.N. Jan, UCSF), UAS-fz1-1 (Bloomington Drosophila Stock Center # 41791) and sevEP-GAL4.B (II) (Bloomington Drosophila Stock Center # 5793).

For experiments with pupae samples, pupae with the correct genotype were collected at 0 hrs after puparium formation (APF) and aged at 28 °C to increase the penetration of genetic perturbations (Das et al., 2002). For experiments with adult samples, flies 3-5 days after eclosion were collected. Both male and female were used for all experiments.

### Heat shock clone induction

Larvae with the correct genotype were heat shocked for 8 - 15 mins at 37 °C at 2 to 4 days after egg laying (AEL).

### Immunohistochemistry and Imaging of Pupae Brains

#### Dissection and Staining

For preparation of pupal brain samples, pupal brains were dissected at the appropriate developmental stages in PBS (Phosphate Buffered Saline) and fixed with 3.7% formaldehyde-PBS for 30 mins. Fixed brains were washed three times in PBT (PBS with 0.4% Triton X-100) at room temperature and then blocked with PBT-BSA (3% Bovine Serum Albumin in PBT) for 1 hour. Two rounds of antibody staining were then performed. In each round, brains were incubated with cocktails of primary antibodies at 4°C overnight, rinsed in PBT, then incubated with cocktails of secondary antibodies at 4°C overnight or for 2 hrs at room temperature, then rinsed again in PBT. For preparation of adult eye samples, eyes were dissected in PBS and fixed with 3.7% formaldehyde-PBS for 30 mins. Fixed samples were washed three times in PBT (PBS with 0.4% Triton X-100) and kept at room temperature in PBT for up to 12 hrs to reduce the pigmentation of eyes. Samples were then incubated with 1:50 Alexa Fluor 633 phalloidin (A22284, ThermoFisher Scientific) for 2 hrs at room temperature and rinsed in PBT.

#### Antibodies

Primary antibodies used for the first round are: mouse anti-Fasciclin II (DSHB, 1D4), 1:20; Chicken anti-GFP (abcam, ab13970), 1:400; Rabbit anti-RFP (Rockland), 1:400. Conjugated secondary antibodies used for the first round are: Goat anti-mouse Alexa-405 (A31553, Thermo Fisher Scientific), 1:250; Goat anti-chicken Alexa-488 (A32931, Thermo Fisher Scientific), 3:500; Donkey anti-rabbit Alexa-568 (A10042, Thermo Fisher Scientific), 1:500. Primary antibody used for the second round is: mouse anti-Chaoptin (DSHB, 24B10), 1:20. Conjugated secondary antibody used for the second round is: Donkey anti-mouse Alexa-647 (A31571, Thermo Fisher Scientific), 1:100.

#### Mounting and Imaging

Samples were mounted in VECTASHIELD Antifade Mounting Medium. Images were obtained on a Nikon A1R-Si inverted confocal microscope with 4 line laser unit (405/488/561/640) and with a 60X oil objective. Z stacks were acquired with a step size of 0.125 μm between optical sections.

### Image Processing for Visual Inspection and Figure Generation

Images were transformed from ND2 format to TIF format and background subtracted in batch using custom ImageJ macro script on a HPC cluster (code available on GitHub).

To calculate the penetration of *sev>Fz* and *sev>N*^*ic*^ perturbation (Table S1), images of *sev>Fz* lamina samples at 38 hrs APF and sev>N^ic^ lamina samples at 45 hrs APF were visually inspected using Fiji (http://fiji.sc/) and targets of bundles with two R3s (bundles with two RFP-positive R cell growth cones and no GFP-positive R cell growth cones) or two R4s (bundles with two GFP-positive R cell growth cones) are counted. Only bundles with R cell growth cones that can be easily traced from origin to target are included in the counting. Fz and Nic perturbation were scored at different developmental time points due to the decay of quality of GFP signal in *sev>Fz* samples over time.

To generate representative images of bundle targeting phenotypes (Figure 1 and Figure 1 – figure supplement 2), we annotated cropped confocal images of *sev>Fz* samples at 38 hrs APF and *sev>N*^*ic*^ samples at 45 hrs APF in Amira 2020.1 (FEI Visualization Sciences Group). For each cropped image stack, growth cones from one bundle were manually segmented in both GFP and RFP channels. Segmented growth cones were then rendered with the appropriate color (red for RFP and green for GFP) in volume, and the unsegmented FasII channel was overlaid as an ortho-slice in gray-scale. TIF files of the results were then exported from Amira and further annotated using Adobe Illustrator. To show bundles in consistent orientations, some images were rotated and/or flipped. Images were also cropped to highlight the representative bundle.

To generate representative images of bundles from 22 to 36 hrs APF (Figure 2, Figure 3A, Figure 3 – figure supplement 1 and Figure 4 – figure supplement 1), images of lamina samples were inspected and adjusted using Fiji (http://fiji.sc/). Brightness and contrast of individual channels were adjusted separately, and only one z-stack was selected for visualization purpose. Images are also cropped to highlight the representative bundle. Tiff files of individual channels were exported from Fiji and further annotated using Adobe Illustrator. To show bundles in consistent orientations, some images were rotated and/or flipped. Images were also cropped to highlight the representative bundle.

### Image Quantification Using Standardized Coordinates

#### Pre-preprocessing

Images of laminas were transformed from ND2 format to TIF format and background subtracted in batch using custom ImageJ macro script on a HPC cluster (code available on GitHub).

#### Annotation

Images were then visually inspected, cropped and annotated using Fiji (http://fiji.sc/). Due to the variation in mounting and sparseness of labeling, only images of the lamina plexus with large intact regions were further analyzed. Images were cropped (in all directions) to keep the part of the lamina plexus that had sparse-enough labeling of growth cones. These cropped images were then used to manually annotate the position of growth cone heels (starting positions) and targets using the Multi-point tool. Heels of growth cones were positioned based on the 24B10 channel while targets were positioned based on the FasII channel. Heels are annotated with points while targets are annotated with ellipses. Multiple z-slices were used to annotate target ellipses to better represent the boundaries of FasII staining. X, Y, positions of heels and major, minor axis lengths and the major axis angle of the target ellipse were exported to a csv file for later quantification. Mapping of each bundle number to its corresponding target numbers was also noted in another csv file. Bundles with rotational defects were not included in the annotation.

#### Quantification

##### Standardized coordinate system

We used custom Python scripts (code available on GitHub) to resample GFP and RFP images of each bundle to obtain representative density maps according to a standardized coordinate. Our standardized coordinate system (see Figure 2) resembles the polar coordinate system. The center of the coordinate system (C = (0,0)) is defined as the intersection of the lines connecting R3 and T3 and R4 and T3’. The polar coordinate is normalized so that: 1) the radius |C-T4| = 1 (A.U.) and 2) the targets T3 and T3’ are placed at angles +1 and −1 (A.U.), respectively. The centers of the target ellipses were used as reference points for the standardized coordinate system.

##### Density map of image slice

We converted image data to a density map in our standardized coordinates in two steps. First, we created a coordinate grid (the radius ranged from 0 to 3.8 with 0.05 intervals; the angles ranged from −3 to 3 with 0.05 intervals). Second, we used the N-dimensional piecewise linear interpolation function within the Python numpy package (v1.16.4) to create a map from Cartesian to polar coordinates; this allowed us to convert GFP and RFP images into density maps in the new coordinate system.

##### Density map of bundle

Density maps for each bundle were computed using the mean density map across z-stacks containing 41 slices, which were centered around the z-slice showing the longest growth cone (typically R3 and/or R4). We manually annotated the length and angle of R3 or R4 growth cones of a given bundle according to the RFP or GFP density map, respectively. Length and angle of growth cones were annotated based on the longest filopodia in the front of the growth cone. Growth cones located in regions where GFP or RFP signals was to dense to distinguish the front were excluded from the annotation. If growth cones exhibit split morphology (i.e., two or more major long filopodia in the front), angles are calculated by the mean of these filopodia. Growth cones were labeled as R3 or R4 based on the absence or presence (respectively) of GFP signals.

### Simulation of Repulsion Model

Heel and target positions were mapped to the standardized coordinate system. Simulated extension angles 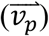 were calculated based on weighted vector sum of two vectors 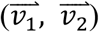. For extension angles of R3, the repulsion vectors: 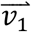 was taken to be the unit vector from the R2 to R3 heels; and 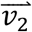 was taken to be the unit vector from the R4 to R3 heels. For extension angles of R4, the repulsion vectors: 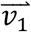 was taken to be the unit vector from the R5 to R4 heels; and 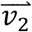 was taken to be the unit vector from the R3 to R4 heels. The weight of each vector represented the strength of its repulsive force. For simple repulsion from neighboring heels, 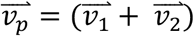. To estimate unequal influence of neighboring growth cone heels, linear regression was performed on 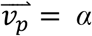 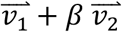 using pooled data from wild-type measurements between 22 to 26 hrs APF. Only data from bundles that were relatively symmetric in shape 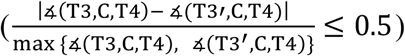 were included in the regression analysis. R3 and R4 angles were fitted independently. Regression analysis was implemented in Python using the scikit-learn package (v0.23.2).

### Statistical Analysis

Sample sizes for each experiment are provided in the figure legends. Statistics were computed in Python using the scipy (v1.2.1) and scikit-posthocs (v0.6.4) packages. A two-sided Mann-Whitney U test was applied when there were only two groups of data being compared. When there were more than two groups, we applied a Kruskal–Wallis H test followed by a post-hoc two-sided Mann-Whitney test with p values adjusted by Bonferroni method. The error bars displayed in all figures represent standard deviation of the mean. Data collection and analysis were not conducted blind to the conditions of the experiments.

## Acknowledgements

We gratefully acknowledge helpful discussions and feedback from Thomas R. Clandinin, Graeme Davis, Marion Langen, and Orion Weiner during the project. We also wish to thank the Clandinin, Hassan, Hiesinger, and Jan labs for fly lines. Finally, we thank Peter Robin Hiesinger, Egemen Agi, and members of the Altschuler-Wu lab and Hiesinger lab for critical comments on the manuscript.

## Author Contributions

Conceptualization and Methodology, W.J., S.J.A. and L.F.W.; Investigation, Formal Analysis, and Visualization, W.J.; Writing – Original Draft, W.J.; Writing – Review & Editing, W.J., S.J.A. and L.F.W.; Supervision, S.J.A. and L.F.W.

---

**Figure 1 – figure supplement 1.**
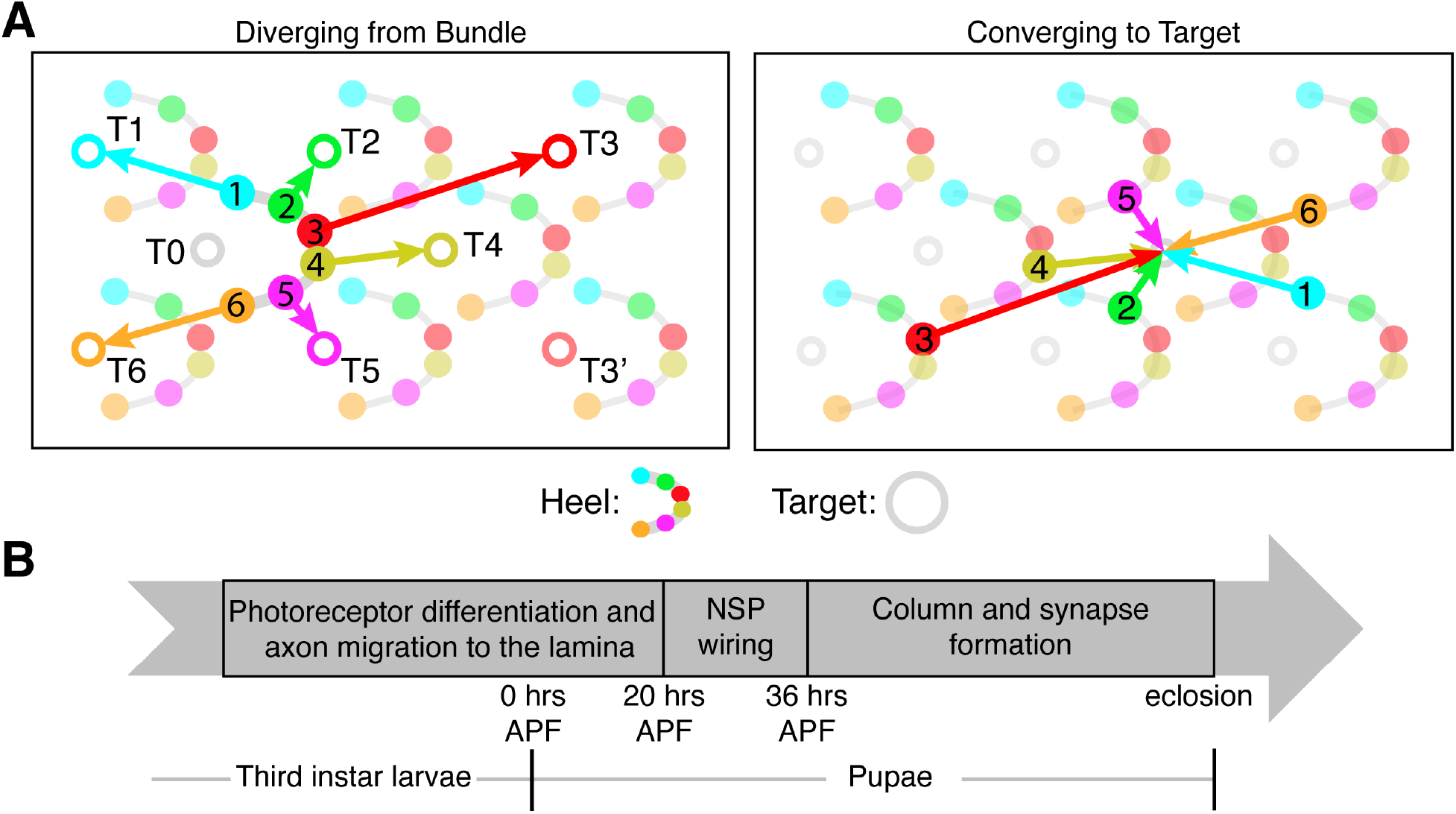
Overview of Neural Superposition. **(A)** Schematic of NSP wiring topology. Solid or open circles: starting points (‘heels”) or targets (respectively). T3’: target of fate-altered R3s; T0: target located within the bundle of interest (though targeted by R cells from other bundles in NSP wiring). (**B)** Schematic for timing of NSP wiring. APF: “After Puparium Formation”.

**Figure 1 – figure supplement 2.**
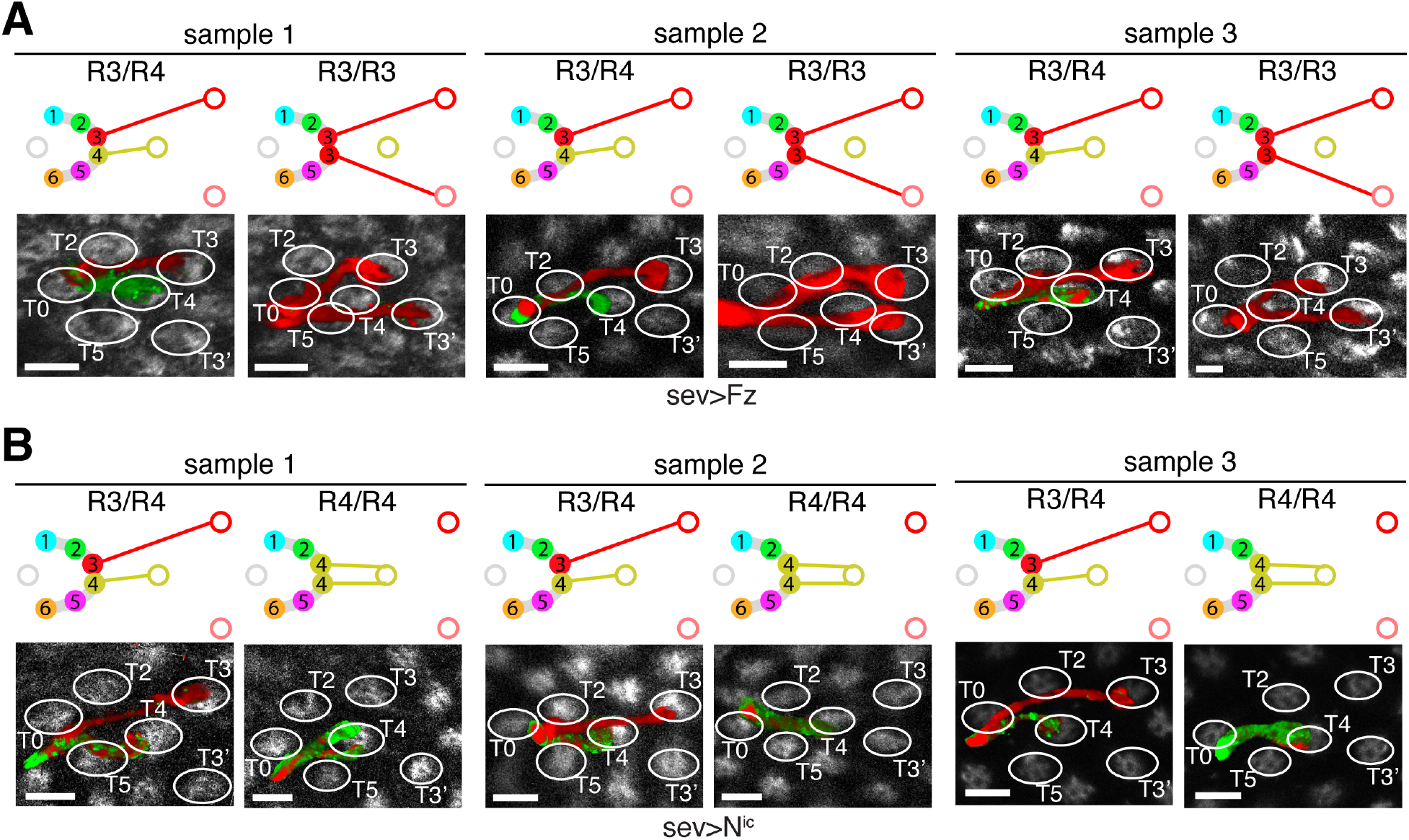
Final targeting of *sev>Fz* and *sev>N*^*ic*^ flies. **(A-B)** Schematics (top panels) and confocal images (bottom panels) of bundles from three **(A)** *sev>Fz* (38 hrs APF) and **(B)** *sev>N*^*ic*^ (45 hrs APF) flies. Top panels: schematic of wild-type or altered wiring topology. Solid or open circles: starting points (‘heels”) or targets (respectively); colors coordinated between R cells and targets. T3’: target of fate-altered R3s; T0: target located within the bundle of interest (though targeted by R cells from other bundles in NSP wiring). Bottom panels: confocal images of representative bundles. Photoreceptor growth cones are segmented and pseudo-colored (Methods) and intensity scaled for visualization. Red: *sev>RFP* expression; green: *mδ-GFP* expression; white: Fasciclin 2 (FasII) antibody staining. White ellipses: targets. Scale bar: 5 μm.

**Figure 3 – figure supplement 1.**
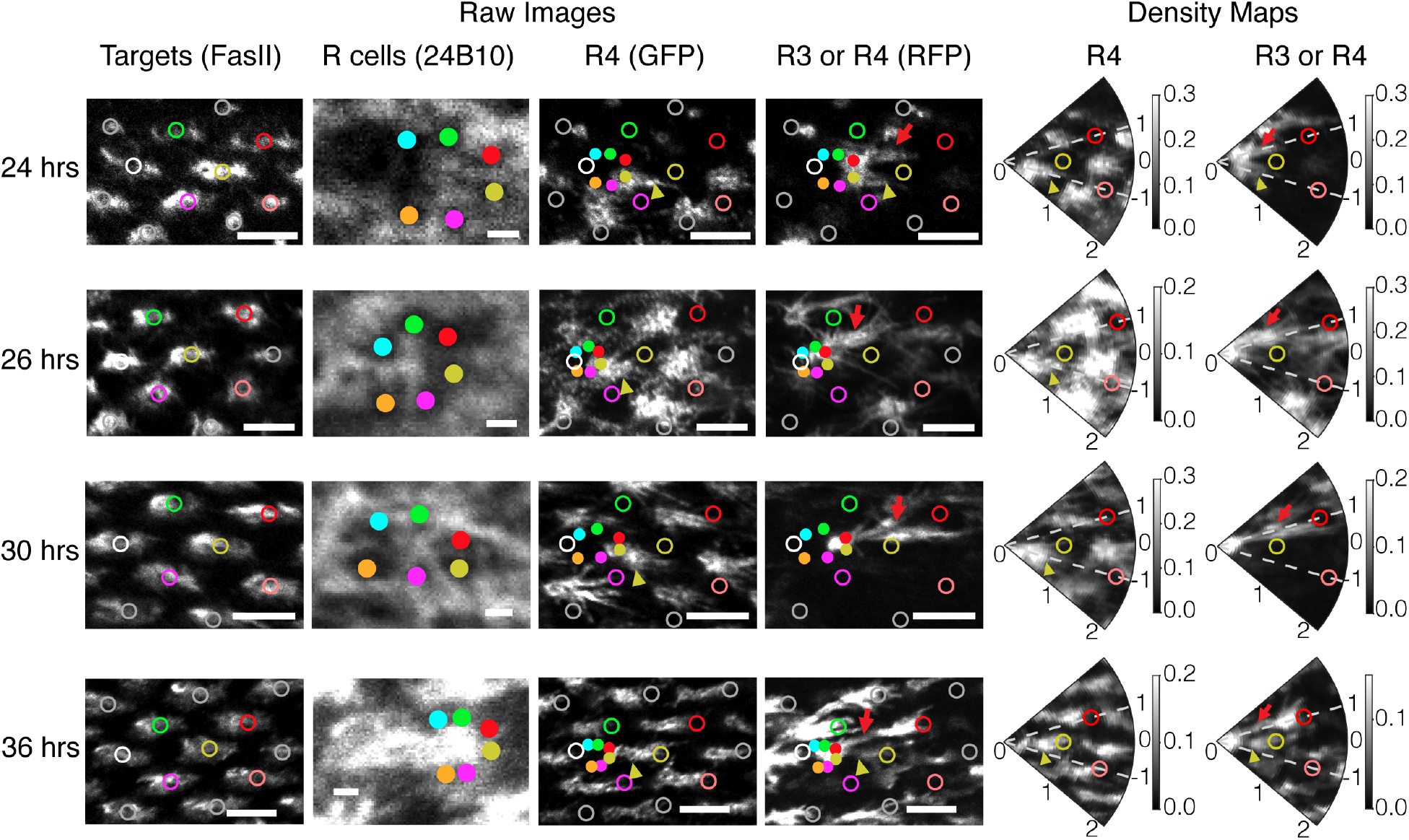
Extension phenotype of wild-type flies over time. Representative images of bundles from wild-type flies at 24, 26, 30, and 36 hrs APF. Left: Raw images of example bundles. From left to right: FasII channel labeling the target cells; 24B10 channel labeling membrane of all R cells; GFP channel labeling membrane of R4 cells; RFP channel labeling membrane of R3 or R4 cells. Right: Density maps of GFP (R4 cells) and RFP channel (R3 or R4 cells) after coordinate transformation. For visualization, intensity is scaled differently for each channel and for each sample. R-cells and targets indicated and colored as in Figure 1 – figure supplement 2; white circles: T0; gray circles: other targets. Yellow arrowheads: R4 growth cones; red arrows: R3 growth cones. Scale bars: 5 μm for FasII, GFP and RFP images; 1 μm for 24B10 images.

**Figure 4 – figure supplement 1.**
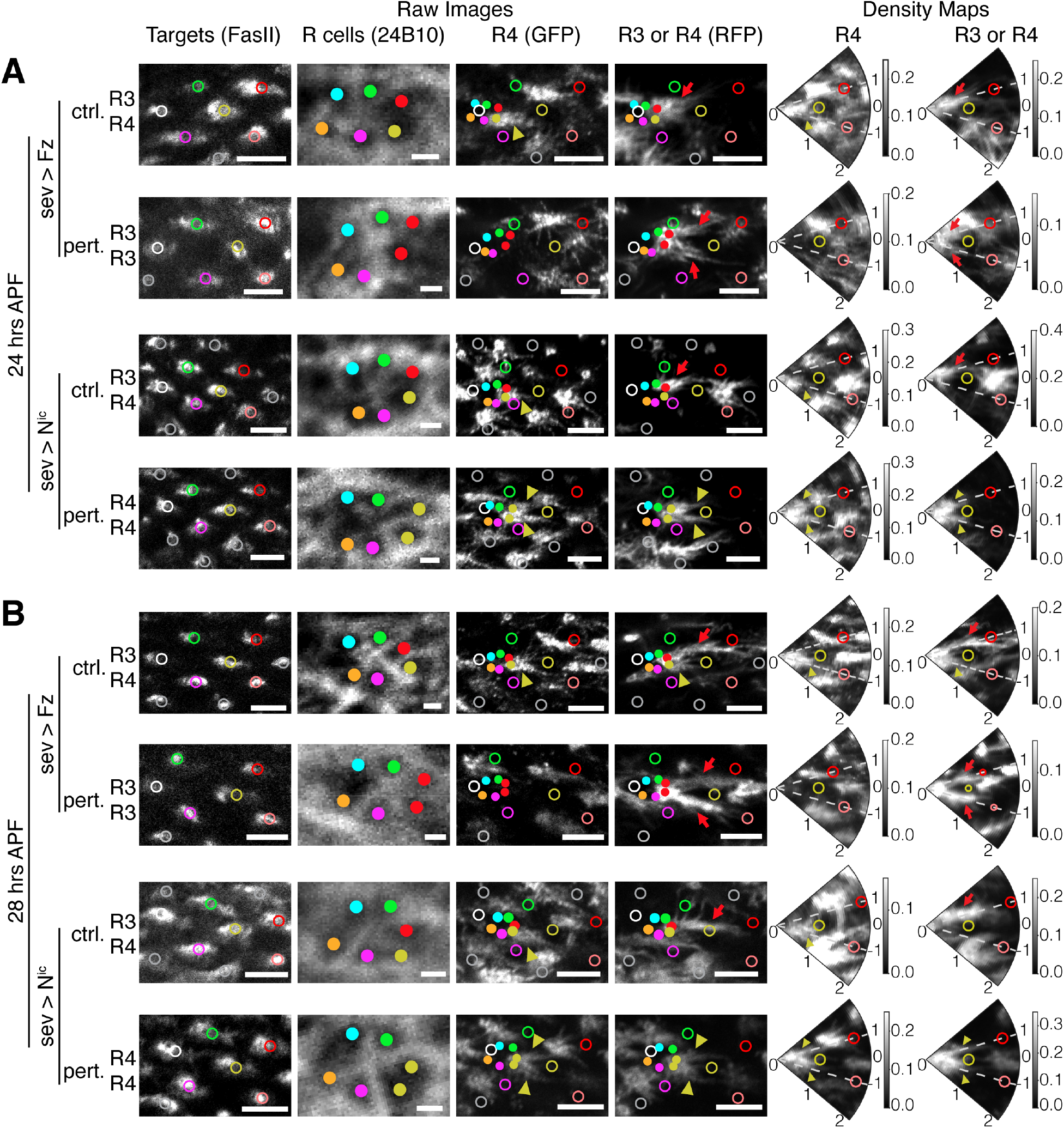
Extension phenotype of *sev>Fz* and *sev>N*^*ic*^ flies over time. **(A-B)** Representative images of wild-type-like (ctrl.) and fate-altered (pert.) bundles in *sev>Fz* and *sev>N*^*ic*^ flies at (**A**) 24 or (**B**) 28 hrs APF. Left four panes are confocal images of example bundles. Right two panels are density maps of GFP (R4 cells) and RFP channel (R3 or R4 cells) after coordinate transformation. Image channels, intensity normalization, annotation and scale bars are as in Figure 3 – figure supplement 1.

**Figure 4 – figure supplement 2.**
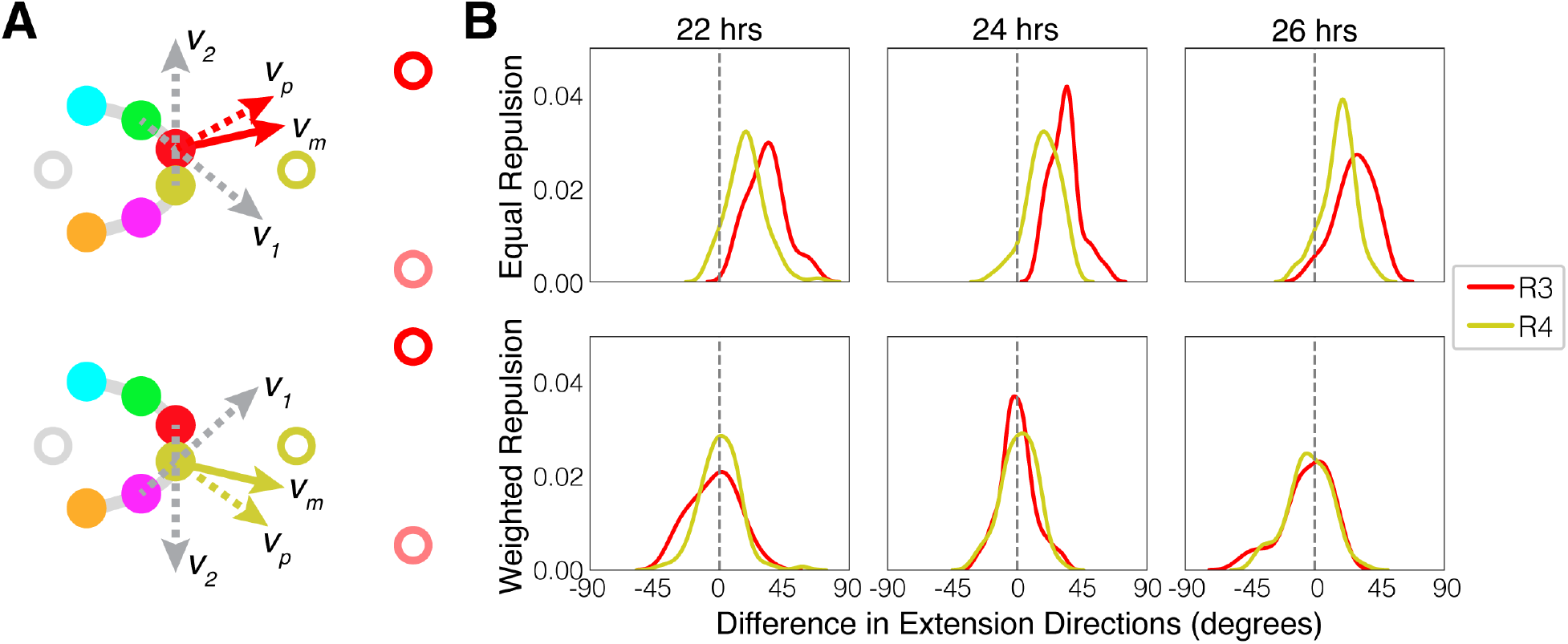
Repulsion model for determining growth cone extension angle. **(A)** Schematics of repulsion model. For R3, 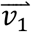 and 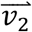 represent repulsive forces from R2 and R4, respectively. For R4, 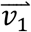 and 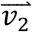 represent repulsive forces from R5 and R3, respectively. 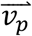: extension direction predicted from simulation; 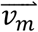: extension direction measured. **(B)** Difference between predicted and actual extension directions for data from 22, 24 or 26 hrs APF. 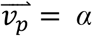 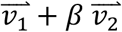 is used to calculate predicted extension directions. For the equal repulsion model, α = β = 0.5. For the weighted repulsion model, linear regression is performed to get α and β that best fit pooled data from wild-type measurements between 22 to 26 hrs APF. R3 regression result: α = 1.03, β = 0.44, R^2^ = 0.75; R4 regression result: α = 1.00, β = 0.64, R^2^ = 0.87.

